# Utilizing single-cell data for per-cell type eQTL mapping in the human pancreas

**DOI:** 10.64898/2026.09.23.753551

**Authors:** Mary Ann Weidekamp, Kim Lorenz, Ruth Elgamal, HPAP Consortium, Klaus H. Kaestner, Kyle Gaulton, Struan F.A. Grant, Benjamin F. Voight

## Abstract

**Aims/hypothesis:** The human pancreas is a central organ for metabolic regulation that is comprised of diverse cell types that uniquely contribute to its function. Previous studies have performed expression quantitative trail loci (eQTL) discovery in either whole pancreas or in pancreatic islets, but due to differences between pancreatic cell types, this approach does not reveal cell type-specific effects. In this study, we sought to either implicate the cell type of action for known eQTLs or identify new eQTLs that may have been masked in bulk studies by performing eQTL discovery in individual pancreatic cell types.

**Methods:** We clustered 153,018 single-cell RNA sequencing (scRNA-seq) data from 71 pancreatic islet donors from the Human Pancreas Analysis Program (HPAP). We performed eQTL discovery in six pancreatic cell types using this resource directly. We further utilized this single cell resource as a reference to deconvolute bulk pancreatic RNA sequencing data from 305 Genotype Tissue Expression (GTEx) project donors and performed eQTL discovery in four pancreatic cell types. Finally, we performed fine-mapping and co-localization of pancreatic cell type eQTLs with metabolic GWAS to connect our findings to metabolic disease risk.

**Results:** From analyzing 71 individuals with single cell profiles, we identified 112 unique eGenes across six pancreatic cell types, 99 of which had been identified previously and 13 unique to this study. From the deconvoluted eQTLs, we identified 3,134 unique eGenes across four pancreatic cell types, 116 of which were unique to our study. Fine-mapping and co-localization of eQTLs with metabolic GWAS yielded key leads that warrant further investigation, such as the association of rs2168101 with *LMO1* expression in alpha cells.

**Conclusions/interpretation:** We identified new signals that were previously not found in bulk pancreatic eQTL studies and potential cell type of action for several signals that were identified previously. Although there are limitations to the power, and therefore, discoverability of this study, it provides insights into how individual pancreatic cells differently contribute to metabolic disease.

**RESEARCH IN CONTEXT:** *What is already known about this subject?:*

- Individual pancreatic cell types have distinct functional roles that differently contribute to metabolic traits and disease risk.
- Expression quantitative trait loci studies (eQTLs) can help connect variants to changes in gene expression, which when paired with association studies, can implicate genes with roles in metabolic disease.
- Previous pancreatic eQTL studies have been carried out in whole pancreatic tissue or whole pancreatic islets, but not at individual cell type resolution.

*What is the key question?:*

- By performing eQTL analyses at individual cell type resolution, can we map known eQTLs to putative cell types of action plus uncover new eQTLs that were previously undetected by bulk pancreatic studies?

*What are the new findings?:*

- Across two eQTL discovery experiments, consisting of single cell data directly and deconvoluted bulk expression profiles, we identified 112 and 3,134 genes associated with an eQTL, respectively.
- Of these results, 13 and 116 of these genes had not previously been associated with pancreatic eQTLs and were therefore unique to our discovery effort.
- After co-localization with metabolic trait GWAS, we observed novel variant-to-gene connections for future follow up, such as rs2168101 for *LMO1*, an eQTL we identified in alpha cells that co-localized with fasting glucose levels.

*How might this impact clinical practice in the foreseeable future?:*

- This study provides new insights into the role of individual cells in metabolic traits, informing future studies on the role of the pancreas in conferring disease susceptibility.

## INTRODUCTION

The pancreas is a central regulator of metabolism, and its dysregulation contributes to the pathogenesis of complex diseases including type 1 and type 2 diabetes, obesity, and pancreatic cancer.[1, 2] Many genetic loci associated with susceptibility for these diseases have been identified genome-wide, and the established physiological role of the pancreas in these diseases implicates it as a key tissue of action for risk variants. The pancreas is composed of distinct cell types, each making unique contributions to its overall function. The endocrine cells of the pancreas, consisting of alpha, beta, delta, epsilon and gamma cells, are located within the islets of Langerhans, and collectively regulate blood glucose through distinct hormonal signaling from each cell type.[3–9] The exocrine compartment, comprising acinar and ductal cells, is responsible for the production and delivery of digestive enzymes to the small intestine.[10, 11] Given the diverse and coordinated functions these cell types perform, placing human genetic associations within specific cellular contexts is critical for identifying the genes and corresponding pathways that modify susceptibility to disease.

A powerful approach to connecting genetic variation to gene expression in specific cellular contexts is through the identification of genetic variation associated with changes in transcript abundance, i.e., expression quantitative trait loci (eQTL).[12–4] When integrated and co-localized with data from complex disease genome-wide association studies (GWAS), eQTLs can pinpoint candidate genes and pathways relevant to disease etiology.[14–6] In the context of the pancreas, existing eQTL resources have been generated largely from bulk tissue or whole islet preparations[17–9]. While generally well powered to associate common variants with expression given the large sample sizes, the resulting datasets obscure the contributions of individual cell types and thus could mask specific signals that are present in distinct cellular populations. In contrast, per-cell-type eQTL discovery can both identify relevant cell types for known signals discovered in bulk or uncover new signals that bulk eQTL analyses fail to detect. Single cell RNA-seq (scRNA-seq) enables cell type-specific expression profiling, and when aggregated into pseudo-bulk profiles, can support per-cell-type eQTL discovery directly, provided the dataset is sufficiently powered.[20]

Deconvolution methods offer an additional, complementary approach that can leverage cell type-specific profiles in the context of bulk data, which often have larger available sample sizes. Previously, expression data across bulk tissues in the Genotype Tissue Expression (GTEx) project v8[19] were deconvoluted using a mouse reference set and the estimated cell type proportions used as covariates to improve eQTL discovery in select tissues.[21] Beyond the application to improve bulk eQTL discovery, deconvolution methods can be applied to perform cell-type eQTL discovery by imputing per-sample per-cell type expression profiles.[22] By leveraging single-cell reference data to decompose existing bulk RNA-seq datasets into their constituent cell type components, this approach extends the utility of well-powered bulk resources.[23–5] Leveraging a bulk resource together with a high-quality human pancreatic single-cell reference dataset makes both single-cell and deconvoluted eQTL discoveries per-cell type tractable in the pancreas.

A recently available scRNA-seq resource generated by the Human Pancreas Analysis Program (HPAP)[26–9] provides an opportunity to conduct per-cell type analysis in the pancreas. This resource has over 150 donors across diverse backgrounds, ages, sexes, and disease status[26]. Multiple assays have been conducted across each donor’s tissue, providing a wide array of data that can be used for analysis. Here, we utilized the HPAP scRNA-seq and whole genome sequencing (WGS) datasets[26] to perform per-cell type eQTL discovery across the major pancreatic cell types. We then employed this single-cell reference to deconvolute bulk pancreatic RNA-seq data from GTEx,[19] enabling eQTL discovery in acinar and ductal cells at greater sample sizes than currently possible with single-cell data alone.

Comparing our cell type-resolved eQTLs to those from prior bulk pancreas and whole islet studies,[17–9] we observed broad concordance of signals while also enabling attribution to likely cell type(s) of action. To understand the disease relevance of these findings, we performed colocalization analyses of cell type-specific eQTLs and GWAS data spanning pancreas-implicated traits including type 1 and type 2 diabetes, obesity, lipid metabolism, glycemic traits, pancreatic cancer, Alzheimer’s disease (sometimes referred to as ‘type 3 diabetes’ due the brain’s relationship with insulin), and sleep.[30–7] These analyses reveal that trait-associated signals are distributed both within individual cell types and shared across multiple cell types, underscoring the value of cell type resolution in interpreting the genetic architecture of pancreas-related disease.

## METHODS

### Data Availability

All data for this project courtesy of the Human Pancreas Analysis Program are available on dbGAP **(Supp Table 1)** and PANC-DB **(Supp Table 1)**[29]. Fastq files for 82 HPAP donors were sourced from PANC-DB (https://hpap.pmacs.upenn.edu). Whole genome sequencing information was sourced from dbGAP **(Supp Table 1)** and included 116 HPAP donors. Five donors that had scRNA-seq data did not have whole genome sequencing data available and were excluded from further analysis **(Supp Figure 1, Supp Table 2)**. Protocols for each individual’s data collection and sequencing are available through PANC-DB **(Supp Table 1)**[29].

**Figure 1.**
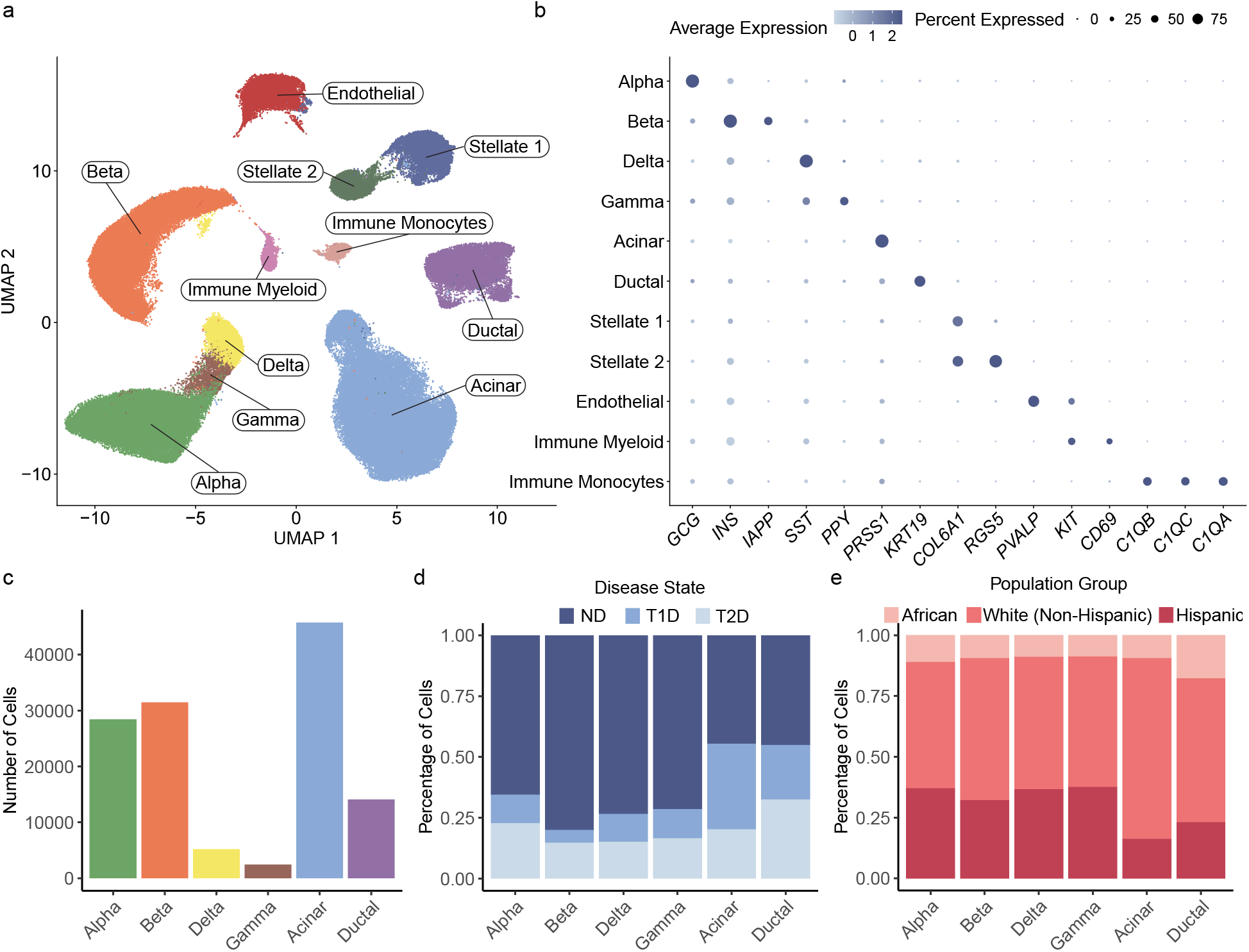
Clustering and Identification of HPAP scRNA-seq data: (a) UMAP plot of the scRNA-seq from 71 HPAP individuals. Each cluster is labeled with the annotated cell type for that cluster based on marker gene identification. (b) Dotplot showing expression of common marker genes in each of the cell type clusters. Size of the dot represents the percentage of cells expressing the marker in that cluster and color of the dot represents the average expression. (c) Number of cells in each of the six pancreatic cell type (alpha, beta, delta, gamma, acinar, and ductal) clusters. (d) Proportion of each cluster that was contributed to by donors with a particular disease status (type 1 diabetic, type 2 diabetic, or nondiabetic). (e) Proportion of each cluster that was contributed to by donors from self-reported population groups (African, White (Non-Hispanic), or Hispanic).

All data for this project courtesy of the Genotype-Tissue Expression (GTEx) consortium are available on dbGAP **(Supp Table 1)** and the GTEx Portal **(Supp Table 1)**[19]. Gene read counts for 328 pancreatic donors from GTEx v8 were sourced from the GTEx Portal (https://gtexportal.org). Whole genome sequencing information was sourced from dbGAP **(Supp Table 1)** and included 305 GTEx donors. 23 donors that had read counts did not have whole genome sequencing data available and were excluded from further analysis.

### Whole Genome Sequencing – Variant Calling and Processing

Data collection for subjects is described elsewhere[29]. Briefly, 30x whole-genome sequencing was performed on 116 HPAP samples at the Broad Institute (2022_08_10 release). A PCR-Free whole genome sequencing (WGS) process was used, which included sample identification quality control (fingerprinting), sample preparation utilizing custom Broad indices (IDT) and Kapa Biosciences HyperPrep library construction. WGS (2x150bp reads) was performed using 2 color chemistry on the NovaSeq platform, with read de-multiplexing, aggregation, and alignment to hg38 with bwa-mem (0.7.15-r1140). Joint variant calling across individuals was performed using standard pipelines (Picard alignment metrics and GATK v3.5).

Subsequently, we assessed and filtered variants across a range of quality control measures using vcftools v0.1.16[38]. These included allele frequencies (freq2), mean depth per individual (depth), mean depth per site (site-mean-depth), per-site SNP quality (site-quality), per-individual missingness (missing-indv), per-site missingness (missing-site), and heterozygosity (het). Based on the results of these analyses, we filtered the whole genome sequencing vcf using vcftools with the following flags: remove-indels, maf 0.01, max-missing 0.75, minQ 30, minDP 15, and maxDP 75. The result was 6,801,817 variants across samples.

### Fastq Processing

Initial processing of the scRNA-seq fastq files was performed using 10x cellranger count version 7.0.0[39] and the reference set Gencode GRCh38 v32/Ensemble 98 (2020-A)[40]. In this workflow, first reads with invalid barcodes are removed. Then, valid barcodes are aligned to the genome (hg18) using STAR[41], with unmapped reads or reads with MAPQ <255 removed from the analysis. After genome alignment, confidently mapped reads undergo transcriptome alignment filtering. In this step, intergenic reads, antisense reads, or reads mapping to more than one gene are removed from analysis. Intronic reads were retained for this analysis, as is standard in cellranger version v7.0 and above. This collection of transcriptomic reads then undergoes unique molecular identifier (UMI) correction and UMI counting. Finally, cell barcodes are called, and a selection of output files are created representing the aligned, filtered, and counted outputs. More details on this method can be found on 10x genomics website **(Supp Table 1).**

### Initial Quality Control

After performing initial processing of the scRNA-seq raw files, additional quality filters were employed to ensure that individuals and data used in downstream analysis are representative of the population. The following quality control steps, including ambient RNA removal, doublet removal, and single-cell clustering protocols are adapted from Elgamal et al, 2023[42]. Upon initial inspection of the cellranger count summary data for number of cells, reads per cell, and genes per cell, two individuals with less than 1,000 genes per cell and four individuals with ambiguous sample labels were removed from downstream analysis **(Supp Figure 1, Supp Table 2)**. The cellranger count outputs from the remaining 71 individuals were read into Seurat v4.3.0[43] using the Read10X function. These individuals’ cells then went through standard Seurat filtering to remove cells with 15% or more reads mapping to mitochondrial DNA, cells with less than 500 UMI reads per cell, or cells containing more than 4,000 UMI reads per cell. The data were then log normalized using a scale factor of 10,000 to ease readability.

### Ambient RNA Removal

When processing cells for scRNAseq, droplets contain some contamination of RNA that does not represent actual transcript levels in the cells. To estimate this contamination and remove ambient RNA, each individual was processed using SoupX (v1.6.2)[44]. This method requires inputs of filtered RNA counts (generated above), raw RNA counts (from cellranger count output), and initial clustering data. Each participant was analyzed through Seurat’s RunPCA, FindNightbors (30 dimensions), FindCLusters (Leiden clustering), and RunUMAP (30 dimensions) to obtain dimension reduction coordinates and clustering estimations that were added as metadata to the count matrices. With these inputs, each participant was analyzed through the automated contamination fraction estimation method from SoupX and their RNA counts were adjusted using the adjustCounts function. After the RNA counts were adjusted, samples were re-corrected for quality control metrics as described above, and all individuals were merged into a single RDS file.

### Doublet Removal

When performing scRNA-seq, some droplets may capture more than one cell, which can confound the results. To retain droplets that contain just one cell (singlets) we used Scrublet (v0.2.3)[45] to predict potential multiplets and remove them from further analysis. Scrublet was run with default settings with the expected doublet rate of 0.06, minimum counts of 2, minimum cells of 3, minimum gene variability percentile of 85%, and 30 principal components. Only cells that were predicted to be singlets were retained for downstream analysis.

### Single Cell Clustering

After quality control, data was clustered to aid in cell type identification using standard Seurat v4.3.0 protocols[43]. Since barcodes were removed in the last step, the data was re-normalized using log normalization at a scale factor of 10,000. The top 2,000 most variable features were selected for clustering using the vst selection method. Covariates were corrected for using 20 principal components and Harmony v0.1.1 batch correction[46]. We specified HPAP donor, tissue source (nPod or Upenn) and 10xGenomics assay chemistry (version 2 or version 3) as Harmony covariates. These metadata are available on PANC-DB[29] **(Supp Table 1).** The data were then piped through runUMAP using harmony reduction with 20 dimensions, findNeighbors using harmony reduction and 20 dimensions and finally the findClusters function with Leiden clustering at a resolution of 0.5. This produced an initial clustering map of the cells across all individuals.

### Cell Type Identification

Once initial clustering is set, it is necessary to identify what cell type is represented by each predicted cluster. We manually annotated our dataset using the distribution of known marker genes for pancreatic cell types [42, 47]. We visualized the distribution of these marker genes using Seurat’s v4.3.0 FeaturePlot and DotPlot functions and compared the top 10 most variable genes for each cluster using the FindAllMarkers[43]. Based on the distribution and enrichment of the identified marker genes, we found that a few clusters showed multiple marker genes but were still separated when checking the distribution in UMAP space. For these cases, the cells from that cluster were selected and re-clustered using Seurat’s v4.3.0 FindClusters function with Leiden clustering and a resolution of 0.25 to create sub-clusters. If visualization and analysis of these sub-clusters using the methods described above could separate identified marker genes, the sub-clustering was kept in the analysis. If one of the sub-clusters contained a high admixture of marker genes that could not be divided, that sub-cluster was removed and the sub-clusters that showed high enrichment of one marker gene were kept. Once any cells in regions that could not be confidently labeled were removed, the dataset was re-run through clustering as described above and final cell labels were assigned based on marker gene distribution.

### CIBERSORTx HiRes Deconvolution

We performed statistical deconvolution of GTEx’s v8 pancreatic RNAseq data using CIBERSORTx v1.0[25]. This was done in two steps, first the fractions mode, then the hires mode. For the fractions mode, we used our HPAP scRNA-seq data with just the non-diabetic individuals as the reference and the GTEx v8 pancreatic RNAseq data[19] as the mixture. We ran fractions mode with S-mode batch correction, the fraction flag set to 0, and single-cell mode set to true. This mode gives us fraction predictions for our data as well as the signature matrix created for our single-cell reference. This signature matrix is a key input for the second step, which is CIBERSORTx HiRes. For this mode, we used the signature matrix as the reference, the same GTEx v8 pancreatic RNAseq data as the mixture and set quantile normalization to true. The per-individual per-cell type reference matrices that are output from this mode were then cleaned to remove genes with NA values (lacked statistical power to impute gene expression) or the value “1” for every individual (indicates there was insufficient evidence of expression).

### eQTL Input file preparation

After clustering or deconvolution, we pulled data for each of our identified cell types to create cell-type specific input files for downstream analysis. We adapted the GTEx v8 eQTL pipeline[19] for our eQTL analysis, treating each cell type as its own tissue. This pipeline requires raw data counts, transcript per million (tpm) counts, and a collapsed gene annotation file as inputs. For each cell type cluster identified above or deconvoluted cell type, the raw count data were utilized to create a pseudo-bulk count matrix per cell type. For the deconvoluted GTEx data, 23 individuals were dropped at this stage that did not have genotype information from the GTEx v8 vcf file[19]. Tpm files for each pseudo-bulk cell type were also generated using a protocol adapted from Elgamal et al[42]. Gene ids in the pseudo-bulk count matrices were compared to gencode release 38, GRCh38.13 comprehensive gene annotation file[40] to collect exons per gene id. Exons were reduced to a set of non-overlapping exons and effective lengths were summed. Tpm values were then calculated by dividing the original counts by the exonic sizes and dividing the result by one million. To create the collapsed gene annotation file, gencode release 38, GRCh38.13 comprehensive gene annotation was uploaded and run through GTEx’s v8 collapse annotation protocol. Once all of the input files were prepared for each cell type, they were individually run through GTEx v8 prepare expression function[19], using default parameters of 0.1 tpm threshold, 6 count threshold, 0.2 sample fraction threshold, and tmm normalization to create input files for eQTL analysis.

### Covariate Analysis

Before calling eQTLs, it is important to prepare a file of covariates for use in the analysis to ensure that external factors that may influence the data can be accounted for. GTEx’s v8 pipeline[19] allows for the creation of a covariates file that combines user specified covariates. For our HPAP single-cell eQTL analysis, we chose factors from each individual’s demographic data (available on PANC-DB): age, sex, and diabetes status. For the GTEx deconvoluted eQTL analysis, we included age and sex as covariate. We also included 10 genotype principal components that help infer factors like population groups. Finally, we calculated 10 peer factors[48] for each cell type input file created above. Each cell type had a final covariates file that included its specific peer factors, the genotype principal components, cohort specific covariates.

### eQTL Discovery

Once the input files for eQTL analysis were completed, we ran the analysis to call the eQTLs. The eQTLs were called using tensorqtl v1.0.8[49], once again treating each cell type as a separate tissue for purposes of our analysis. For each cell type, tensorqtl was run in two different modes. The first was the cis-qtl mapping mode that generates summary statistics across all variant and gene pairs for each chromosome. This outputs a file with data for all pairs tested in the study for use in downstream analysis. The second mode that tensorqtl was run with is the cis-qtl permutation analysis, which is the main mode for tensorqtl and generates phenotype-level summary statistics with empirical p-values, enabling calculation of genome-wide false discovery rate (FDR)[49]. Both of these modes were run with standard parameters with the exception of adding a minor allele frequency threshold of 0.05. From these files, we could calculate significant variant–gene pairs by filtering for pairs with a q-value below 0.05 (below 0.1 for the single-cell eQTLs due to power constraints) and a p-value that was below the nominal p-value threshold for that gene as calculated by tensorqtl. These results were collected into a significant variant gene pairs file.

### eQTL Co-Localization Analysis

To compare eQTLs across cell types within our own study and across previous databases, we performed statistical colocalization using the coloc R package v5.2.3[50]. Each co-localization was done pairwise between two datasets at a time. For each dataset, a list of significant eGenes was obtained and overlapped to retain a list of significant eGenes that were shared. Then, for each shared eGene, all variants tested at the locus were collected from each database and the slope, slope standard error, variant id, position of variant, minor allele frequency, and nominal p-value were curated to run in coloc.abf under trait type quantitative. For the TIGER dataset, the minor allele frequencies were assigned to each variant reported using 1000 genomes reference panel[51]. TIGER variants also were run in coloc.abf using variant id, position of variant, an sdY of 1, the imputed minor allele frequencies, and nominal p-values. Once coloc was run on each shared eGene between datasets, the results were filtered to retain only eQTLs that had a posterior probability of hypothesis four (PPH4) greater than 0.8.

### Open Chromatin Data

For many of our top eQTLs, we wanted to ensure that our eVariants of interest were in open chromatin regions for the cell type they were implicated in. To this end, we used data generated from a previous study[52]. Data were generated using 3 individuals from the HPAP project over all pancreatic cell types and peaks were called as described previously[52]. Data was visualized using the IGV genome browser and was lifted over from GRCh37 to GRCh38.

### ABF fine-mapping

To better pinpoint the most likely causal variants for our eQTL data, we performed statistical fine-mapping using the approximate bayes factor (ABF) framework[53]. For every cell type, significant eGenes and associated genomic regions were tested. The genomic regions used to test each eGene was determined by taking the start point of the furthest upstream variant tested for that gene to the end point of the furthest downstream variant tested for that gene. Each gene and associated region was then run individually using the Wakefield ABF for fine-mapping.

### GWAS Colocalization Analysis

To determine if our identified eQTLs were related to any traits, we performed statistical colocalization using the ColocQuiaL pipeline[14] for eQTL colocalization with GWAS, which utilizes the underlying engine of coloc[54]. The LD reference panels and recombination rate references were obtained from the 1000 Genomes Project using the CEU through IBS populations[55]. The allpairs and significant pairs files were tab indexed, and the tissue file was created to instead hold the names of the individual cell types analyzed, treating each pseudo-bulk cell type result as its own tissue for the purposes of our study. We performed colocalization between eQTL data sets and datasets for type 2 diabetes, type 1 diabetes, BMI, waist-hip-ratio (WHR), WHR adjusted for BMI (WHRadjBMI), HbA1c, LDL, HDL, Triglycerides, Total Cholesterol, Pancreatic Cancer, Insomnia, Sleep Duration, and Alzheimer’s Disease[30–7]. Colocalizations were deemed significant if PPH4 had a value above 0.8.

## RESULTS

### HPAP Dataset Information and Processing

We assembled a diverse cohort of individuals collected and assayed by HPAP. Specifically, this resource consisted of scRNA-seq and WGS data collected from 82 and 116 post-mortem pancreases, respectively. After filtering for individual donors with both scRNA-seq and WGS data available and removing donors that had low-quality sequencing or ambiguous sample labels, we retained 71 individuals for this analysis **(Supp Fig 1a)**. These subjects were distributed across self-reported population groups (44 White (Non-Hispanic), 14 Hispanic, and 13 African American), biological sexes (35 males, 36 females), and diabetes status (43 non-diabetes, 11 type 1 diabetes, 17 type 2 diabetes) **(Supp Table 2)**.

Our cohort of 71 individuals initially yielded 316,957 cells which were subject to stringent quality filters, including removing mitochondrial DNA, filtering for high or low RNA reads per cell, correcting for ambient RNA, and removing predicted multiplets **(Supp Table 1, Supp Fig 1b) (Methods)**. Any cells that could not be confidently labeled in our clustering were also removed to ensure we retained only high-quality populations of each cell type. This resulted in a total of 153,018 cells that were subsequently utilized for downstream analyses **(Supp Fig 1b)**.

### HPAP Cell Type Clustering and Identification

To classify and label our quality-controlled collection of cells, we performed cell type annotation of clusters using the expression of canonical marker genes **(Methods)**. We identified 11 unique clusters, which included populations of pancreatic endocrine alpha, beta, delta, and gamma cells **(Fig 1a)**. In addition, we identified populations of exocrine acinar and ductal cells as well as immune, endothelial, and other cell types **(Fig 1a)**. Each labeled cluster showed enrichment of previously reported marker genes for the associated cell type (i.e., *INS* for beta cells, *GCG* for alpha cells, *SST* for delta cells, *PPY* for gamma cells, *KRT19* for ductal cells, and *PRSS1* for acinar cells)[42, 47] while all other respective non-cell type markers were expressed at low levels **(Fig 1b and Supp Fig 2)**, confirming cluster labeling and purity.

**Figure 2.**
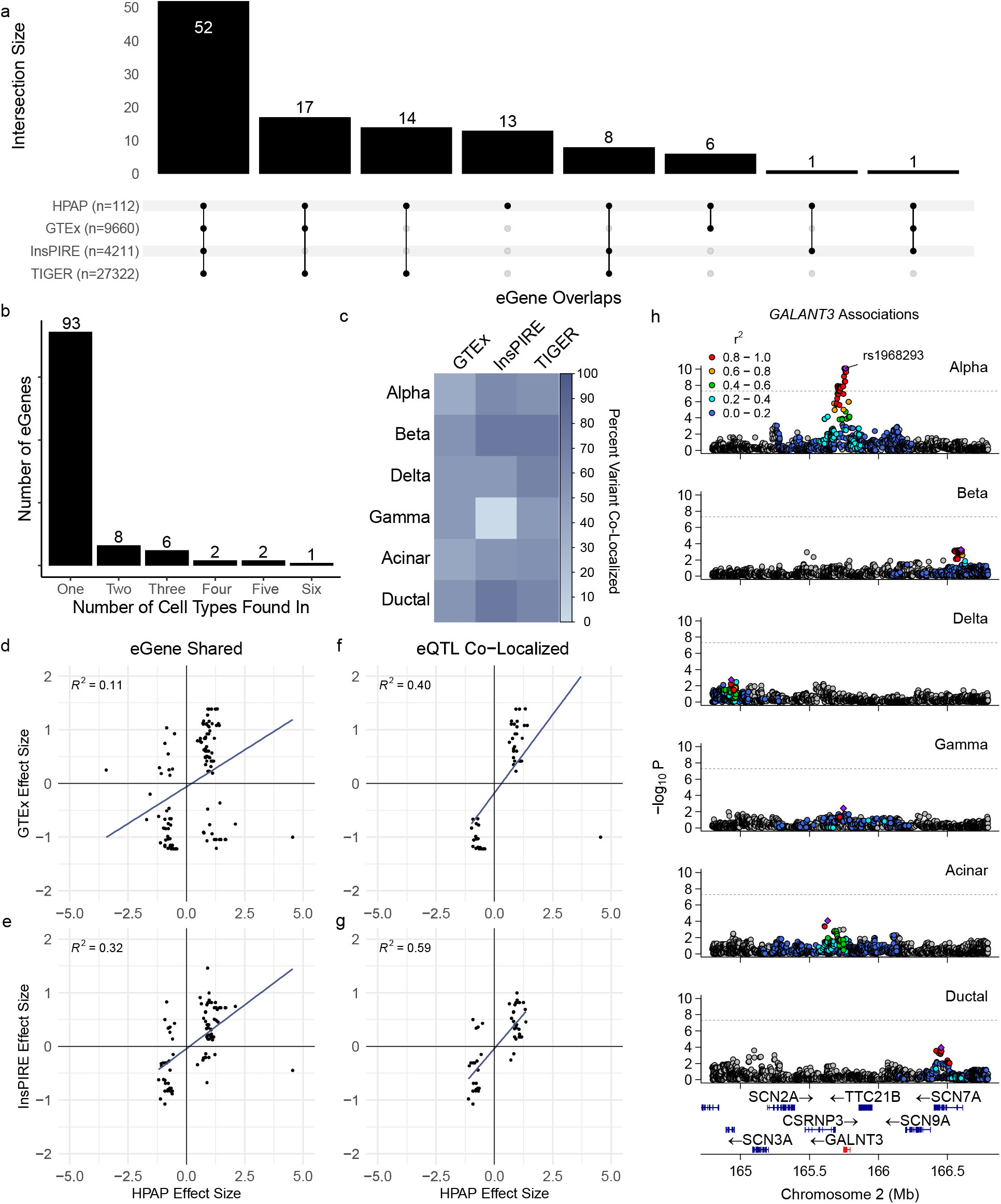
eQTL Discovery from HPAP single Cell data: (a) Upset plot displaying the overlap between the single cell eGenes from HPAP and previous pancreatic eQTL studies. (b) Plot displaying which eGenes were found in just one, or across multiple cell types in the single cell eQTL analysis. A majority of eGenes were discovered in a single cell type. (c) Heatmap displaying the percent of eQTLs that were co-localized between each cell type’s eQTLs and previous pancreatic eQTL studies. This was calculated by taking the total number of eGenes that overlapped and determining how many of them had lead variants for the eQTL that co-localized. (d) Direction of effect comparison for eQTLs that shared an eGene between the single cell eQTLs from HPAP and the GTEx pancreas eQTLs. (e) Direction of effect comparison for eQTLs that shared an eGene between the single cell eQTLs from HPAP and the InsPIRE islet eQTL study. (f) Direction of effect comparison for eQTLs that were fully co-localized between the single cell eQTLs from HPAP and GTEx pancreas eQTLs. (g) Direction of effect comparison for eQTLs that were fully co-localized between the single cell eQTLs from HPAP and the InsPIRE islet eQTL study. (h) Locuszoom plot of *GALNT3* across the HPAP single cell eQTL data. Color indicates the LD score between the displayed variant and lead variant.

We next characterized the frequency of different cell type clusters across subjects. The acinar cluster had the largest number of cells (n=45,745) **(Fig 1c, Supp Table 3)**, followed by beta (n=31,420), alpha (n=28,417), ductal (n=14,059), delta (n=5,181 cells), and gamma (n=2,389) cells **(Fig 1c, Supp Table 3)**. We excluded donors contributing less than 5 cells to a given cluster, resulting in 70 donors for acinar, alpha, and beta cells, 67 donors for ductal cells, 66 donors for delta cells, and 60 donors for gamma cells **(Supp Table 3).** We measured the proportion of cell types by disease state **(Fig 1d)** and population groups **(Fig 1e),** noting qualitative trends in these data, including the expected depletion of type 1 diabetes donors in the beta cell population **(Fig 1d)**, which led us to include these metrics as covariates in downstream analyses.

### HPAP Single Cell eQTL Mapping

To identify genetic variants associated with variation in expression across the major pancreatic cell types alpha, beta, delta, gamma, acinar, and ductal, we next performed eQTL discovery analyses directly using HPAP single-cell data. We aggregated cellular profiles for each cell type into pseudo-bulk populations per donor and performed cell type eQTL analyses using the GTEx analytic pipeline[19] **(Methods)**. Overall, we observed 19,822 eVariants associated with gene expression levels across cell types (FDR < 0.1, Methods). Per cell type, there were 4,259 eVariants for acinar cells, 4,576 for alpha cells, 3,573 for beta cells, 1,848 for delta cells, 3,826 for ductal cells, and 1,740 for gamma cells **(Supp Table 4)**. From these variant gene pairs, we found 112 unique genes with a significant eQTL (eGene) across all cell types **(Fig 2a)**, including 55 acinar, 56 alpha, 14 beta, 5 delta, 19 ductal, and 2 gamma eGenes **(Supp Table 4)**. We found that significant eVariants were localized in open chromatin regions characterized across cell types from HPAP (**Supp Table 4)** and qualitatively were more significant the closer they were to the gene transcription start site (**Supp Fig 3**). These metrics provide additional confidence in the discovered eQTLs.

**Figure 3.**
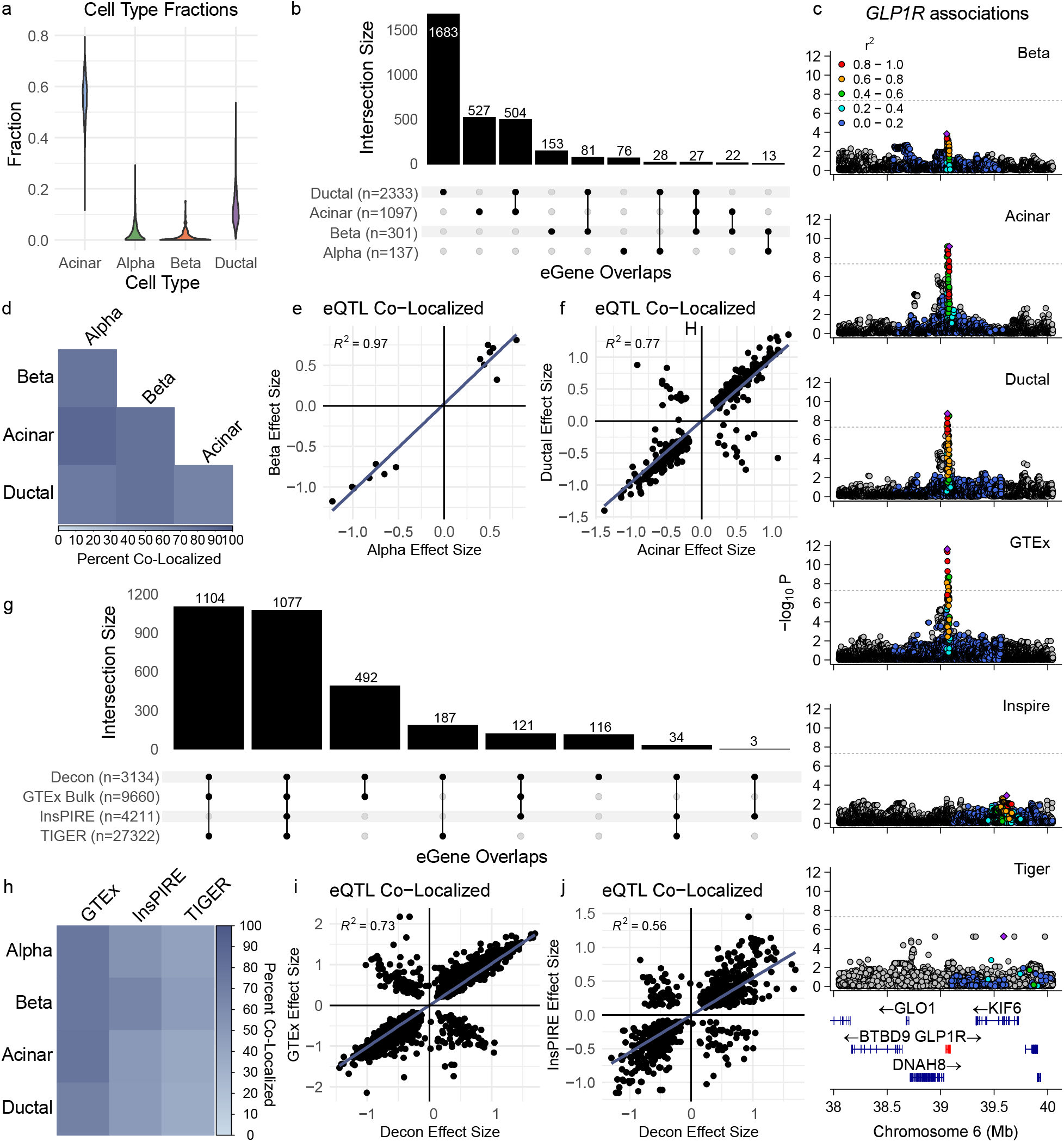
Deconvolution and Individual Cell Type eQTL Discovery of Bulk GTEx v8 Pancreatic RNAseq Data: (a) Violin plot displaying the fractions of each pancreatic cell type detected (Acinar, Alpha, Beta, Ductal) from deconvoluting GTEx v8 bulk pancreatic RNA sequencing data. (b) Upset plot displaying overlap of eGenes across cell types for the deconvoluted eQTL discovery. (c) Locuszoom plot of *GLP1R* across the deconvoluted eQTL cell types and previous pancreatic eQTL studies. There were significant eQTLs detected in acinar and ductal cells as well as the GTEx pancreas eQTL study. (d) Heatmap displaying the percentage of eQTLs that are fully co-localized between the different deconvolution eQTL cell types. Only eQTLs that shared an eGene were tested. (e) Direction of effect comparison between co-localized beta cell eQTLs and alpha cell eQTLs. (f) Direction of effect comparison between co-localized acinar and ductal cell eQTLs. (g) Upset plot showing overlap of the eGenes from the deconvoluted eQTL study and previous pancreatic eQTL studies. (h) Percent of eQTLs fully co-localized between the cell type specific deconvoluted eQTLs and previous eQTLs studies. Only eQTLs that shared an eGene were tested between groups. (i) Direction of effect comparison for eQTLs that fully co-localized between the deconvolution study and GTEx pancreatic eQTL study. (j) Direction of effect comparison for eQTLs that fully co-localized between the deconvolution study and the InsPIRE islet eQTL study.

We next determined the degree to which eQTLs were shared between pancreatic cell types. We found that most eGenes were found in just one of our cell types, and the number of eGenes shared continued to decrease as more cell types were compared **(Fig 2b, Supp Figure 4).** For any eQTL that did have an eGene shared between one or more cell types, we co-localized the eVariants to further determine whether there was a shared eQTL. We found that 63% of the eQTLs that shared an eGene also had a co-localized eVariant, and all co-localized eVariants had the same direction of effect **(Supp Table 5)**.

**Figure 4.**
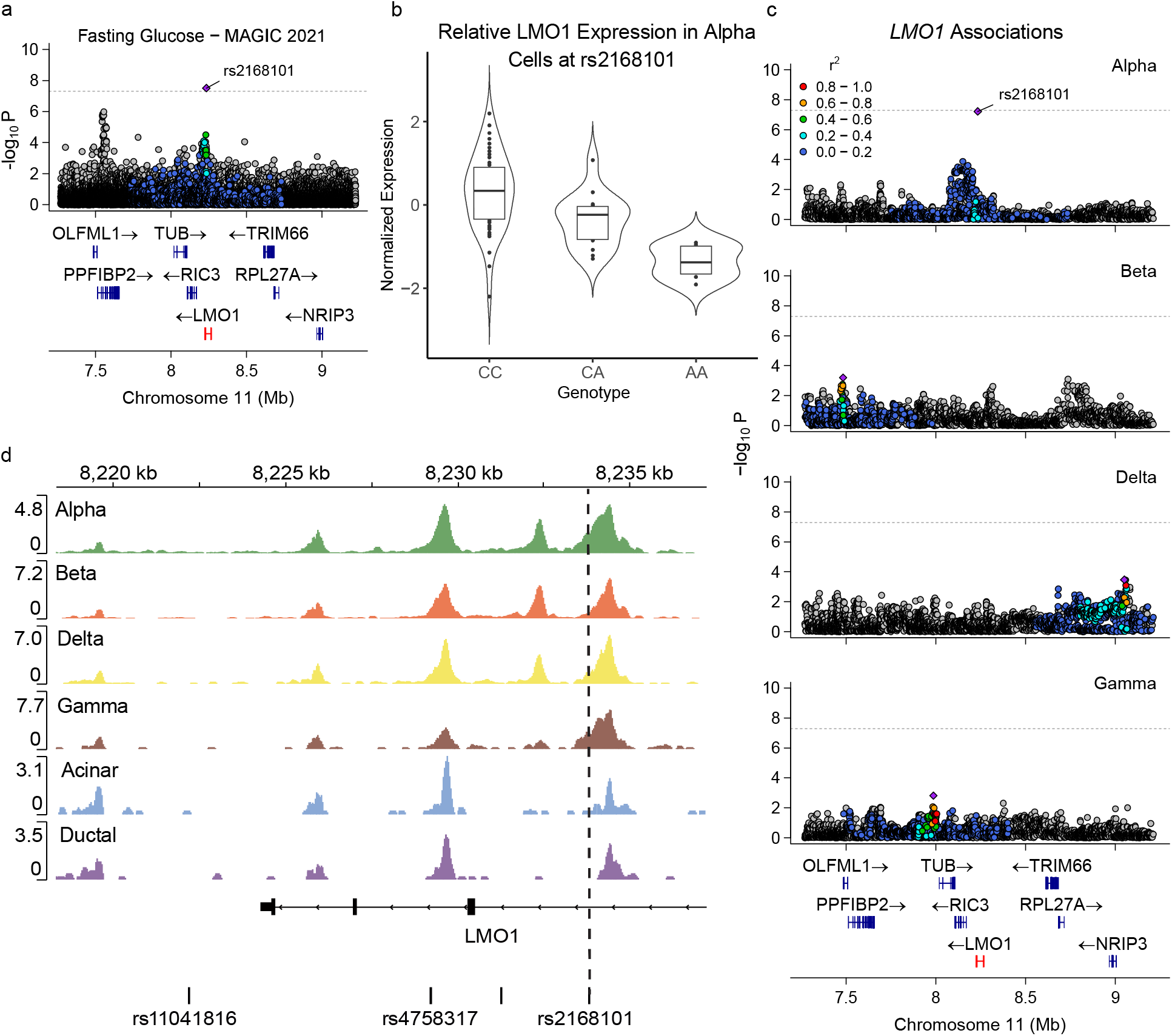
Characterization of rs2168101 and LMO1: (a) Locuszoom plot of rs2168101 for the MAGIC consortium’s fasting glucose GWAS study. Color indicates the LD between the displayed variant and lead variant. (b) Violin plot of LMO1 expression in alpha cells identified from the HPAP single cell study relative to the variant rs2168101. (c) Locuszoom plot of the LMO1 locus across the endocrine cell types from the HPAP single cell eQTL analysis. Color indicates LD between the displayed variant and lead variant. (d) Regions of open chromatin for each HPAP cell type as described in Su et al. Dashed line shows location of rs2168101.

### Comparison of HPAP eQTLs to Previous Studies

To contrast our single cell-derived study results with previous bulk studies, we compared eQTL results eQTLs in whole pancreas from GTEx (n=305, >9.6k eGenes)[19], eQTLs in pancreatic islets from InsPIRE (n=420, >4.2k eGenes)[18], and eQTLs in pancreatic islets from TIGER (n=404, >27k eGenes)[17]. At the eGene level, we observed that 52 of the 112 eGenes detected were consistently observed across all three sets (GTEx, InsPIRE, and TIGER) (**Fig 2a)**. Almost every eGene detected in our study was found in at least one other database, while 13 eGenes were unique to our single cell eQTL discovery effort **(Fig 2a)**.

For the eGenes that were shared between studies, we assessed the degree to which eQTLs were shared through co-localizing the eVariants and assessing the effect direction. We found that for eQTLs that shared an eGene, 54% of the eVariants were co-localized across studies **(Supp Table 6)**. Per cell type, we observed the highest co-localization with beta cell eQTLs (between 54% and 75%), with the lowest agreement against GTEx **(Fig 2c)**. This was unsurprising given that the GTEx data would be made up primarily of exocrine cells. We directly compared the direction of effect between our discovered eQTLs and GTEx or InsPIRE databases, given they reported both effect size and direction[18, 19]. We compared the direction of effect in two different ways, first by comparing top eQTLs that just shared an eGene and then comparing eQTLs whose eVariants were also co-localized. We found a higher level of agreement when comparing effect direction for eQTLs that co-localized on the variant level **(Fig 2d-g).** When we compared top eQTLs that shared an eGene, only 75-80% of the eQTLs shared a consistent direction of effect **(Fig 2d, e).** However, when we assessed those with a co-localized eVariant as well, this increased to between 88-97% of eQTLs sharing a direction of effect **(Fig 2f and 2g).** Taken together, these results reveal the importance of variant-level comparisons when evaluating whether eQTLs are shared.

One of our goals in comparing our study to previous pancreatic eQTL databases was to uncover cell-type specific eQTLs by generating evidence supporting the cell type of action for a known eQTL. One example is the eQTL for *GALNT3* at rs1968293. In our study, a significant eQTL for *GALNT3* was found only in alpha cells **(Fig 2h)**. However, InsPIRE previously detected a significant eQTL for *GALNT3*, and we found that their lead eVariant (rs2304002) co-localized with our alpha cell variant **(Supp Fig 5, Supp Table 6)**. While this result provides evidence for alpha cells being cell type of action for the *GALNT3* eQTL, it is possible more cell types could be implicated with a higher powered experiment. We also sought to detect previously undiscovered eQTLs that could have been masked in rarer cell types when evaluating bulk data. Indeed, we identified *HEATR5B* as an eQTL found only in our alpha cell population **(Supp Fig 6a)**, while in contrast no significant eQTL was detected for *HEATR5B* in any of the bulk pancreatic eQTL databases **(Supp Fig 6b)**.

### Deconvolution of Bulk RNAseq Data

While our single cell dataset has clear limitations in power for eQTL discovery at current sample sizes, it still represents a catalog of cell type expression profiles that can be used in deconvolution analyses of bulk pancreatic data. Towards that end, we utilized the scRNA-seq data available from HPAP from our previous experiments as a reference set for deconvolution of GTEx’s bulk pancreatic RNA-seq data. The GTEx dataset is comprised of 328 donors, 305 of which had WGS data available[19]. These data were generated from whole pancreatic tissue, where exocrine acinar and ductal cells represent the majority of cells with only modest populations of the endocrine cell types. By using deconvolution, we sought to generate per-cell type expression data for these cells to allow for a higher-powered cell type specific eQTL discovery effort.

We performed deconvolution analyses on the GTEx v8 pancreatic data using the CIBERSORTx HiRes pipeline. We leveraged all non-diabetic HPAP donors and ran CIBERSORTx as described **(Methods)**. We found that acinar cells were the most abundant cell type overall, with an across subjects median of 55.93% of cells detected **(Fig 3a, Supp Table 7).** The next most prevalent cell type was ductal cells, with a median proportion of 11.57% **(Fig 3a, Supp Table 7)**. These results were expected, given that the GTEx bulk pancreatic RNA-seq data was generated from whole pancreas, primarily consisting of exocrine cells[56]. As for the endocrine pancreatic cell populations, we detected median values of 1.31% for alpha cells, 2.48% for beta cells **(Fig 3a)**, less than 0.1% for delta cells, and 6.86% for gamma cells. The median calculated Pearson correlation across individuals was 0.8479 and the median root mean squared error was 1.0374 **(Supp Table 7)**. The high gamma cell proportion was unexpected, as they are a relatively rare cell type in the pancreas[57]. Given this, we elected not to proceed with eQTL analysis of delta cells, which were detected at very low rates, or gamma cells, which were likely misclassified.

### eQTL discovery of deconvoluted GTEx data

After deconvoluting GTEx v8 pancreatic bulk RNA-seq data and generating per-individual, per-cell type expression matrices for the acinar, ductal, alpha, and beta cells, we next performed eQTL discovery on these data. We first filtered our expression matrices by retaining only individuals with at least a 0.5% portion of the cell type we were analyzing. This led to the retention of all 305 GTEx donors for acinar and ductal cells, 185 donors for alpha cells, and 139 donors for beta cells **(Supp Table 8)**. We next selected genes for testing analogous to the filtering parameters set in our earlier eQTL analysis **(Methods)**. This led to the retention of 14,852 genes for testing in acinar cells, 16,361 in ductal cells, 11,437 in beta cells, and 6,830 in alpha cells **(Supp Table 8).** We performed eQTL discovery analysis, separately for each cell type, using the WGS from GTEx, and including 10 genotype principal components, 10 peer factors, age, and sex as covariates **(Methods)**.

From this analysis, we identified 3,134 unique eGenes across our four pancreatic cell types **(Fig 3b)**. We observed the most eQTLs in our ductal population, with 326,395 eVariants associated with change in expression across 2,333 eGenes **(Fig 3b, Supp Table 8).** This was followed by acinar cells with 163,746 eVariants in 1,097 eGenes, beta cells with 44,621 eVariants across 301 eGenes, and alpha cells with 13,631 eVariants across 137 eGenes **(Fig 3b, Supp Table 8).** As with our HPAP eQTL analyses, we assessed if the eVariants we observed were found in regions of open chromatin in their respective cell types and found that in all cell types tested and found that >98% of the eVariants were found in regions of open chromatin **(Supp Table 8).** We observed that, qualitatively, eVariants with the most significant *P*-value were on average closer to the TSS for their given eGene **(Supp Fig 9).** Additionally, the cell types that were more abundant in the GTEx data and that had more individuals and genes to test had higher eQTL discovery numbers, highlighting that the power to detect islet eQTLs remained limited in this context.

We next assessed the extent to which eQTLs discovered via deconvolution were shared across cell types. At the level of eGenes, we found that a majority were detected in just one cell type, but there were more overlaps than what we observed in our single cell eQTLs. Across all four cell types, approximately half of the eGenes were detected in just that cell type, while the other half were shared with one or more other cell type(s) **(Fig 3b)**. Acinar and ductal cells had the most overlap with 504 eGenes shared, which was just under half of all acinar eGenes we discovered and 21% of ductal eGenes **(Fig 3b)**. One of these shared eGenes was *GLP1R,* which was shared between acinar and ductal cells, but not in beta cells **(Fig 3c)** or alpha cells. At the variant level, we tested for co-localization of the eVariants for any shared eGenes between cell types to assess if the full eQTL was shared. We found that eVariants had a high percentage of co-localization, between 67 and 88% **(Fig 3d, Supp Table 9)**. For eQTLs that fully co-localized across cell types, we tested the direction of effect, finding that there was a high correlation across cell types **(Fig 3e and 3f, Supp Fig 10).** Our example of *GLP1R* had co-localized eVariants between acinar and ductal cells with a consistent effect direction **(Supp Table 9).** Overall, we observed a high level of agreement across cell types, especially between the two exocrine cell types, but there remained additional signals detected in just one cell type.

### Comparison of deconvolution eQTLs with previous studies

After generating a catalog of eQTLs from deconvoluted data, we tested for signals found specifically in this analysis that were undetected in other bulk studies. We thus compared deconvoluted eQTLs with those from the GTEx v8 pancreas eQTL analysis, the InsPIRE dataset, and the TIGER dataset. Among our 3,134 unique deconvoluted eGenes, only 116 were not detected in previous studies **(Fig 3g)**. This is consistent with the fact that this comparison includes bulk GTEx eQTL analysis, which analyzed the same RNA-seq and WGS data, albeit in bulk. Indeed, 2,794 of the eGenes were shared between the bulk GTEx analysis and the deconvoluted eQTLs **(Fig 3g).** One of the shared hits between bulk GTEx and the deconvoluted eQTLs was *GLP1R*, which was not significant in either the InsPIRE or Tiger studies **(Fig 3c)**. Of the hits that were unique to this study, a majority resulted from the ductal cell population, with 71 of the 116 unique eGenes being found in the ductal cell population **(Supp Fig 11).** This far outweighed the other cell types, with 10 of the unique eGenes being found in alpha, 9 in beta, and 27 in acinar cells **(Supp Fig 11).**

We also compared the deconvoluted data to previous studies at the level of eVariants to assess the extent to which full eQTLs were shared across databases. We found that eGenes that overlapped with GTEx bulk eQTL analysis also had the highest levels of variant co-localization, with between 68-77% co-localizing across our cell types **(Fig 3h, Supp Table 10)**. We observed much lower agreement with InsPIRE and TIGER’s eQTLs, with only between 36-65% co-localizing with these islet-focused studies **(Fig 3h, Supp Table 10).** When comparing the direction of effect for these co-localized eQTLs, we also found a high level of agreement with GTEx’s bulk eQTL analysis, with 2,338 eQTLs having the same direction of effect and only 204 yielding an opposing direction of effect **(Fig 3i)**. The *GLP1R* eQTL in both acinar and ductal cells co-localized with the significant GTEx variant and had the same direction of effect **(Supp Table 10).** We observed similar patterns with eQTLs that co-localized with InsPIRE, where 706 had the same direction of effect while 96 opposed **(Fig 3j)**.

### Approximate Bayes Factor fine-mapping of eVariants

After characterizing eQTLs from both our single cell and deconvolution eQTL discovery efforts, we next sought to prioritize potential variants of action for each signal through statistical fine-mapping. To this end, we calculated approximate bayes factor (ABF) analyses at each significant eQTL signal to identify the credible set and likely causal variant **(Methods)**. We did this for both the single-cell and deconvoluted eQTLs. For our single-cell eQTLs, we derived credible sets for 55 alpha eGenes, 13 beta eGenes, 5 delta eGenes, 2 gamma eGenes, 53 acinar eGenes, and 18 ductal eGenes **(Supp Tables 11-16)**. For our deconvoluted eQTLs, we derived credible sets for 137 alpha eGenes, 301 beta eGenes, 1,085 acinar eGenes, and 2,291 ductal eGenes **(Supp Tables 17-20)**.

### eQTL co-localization with related GWAS

We next sought to connect eQTL signals from our study with human traits and diseases previously implicated in pancreatic function[58–3] to predict the genes involved in disease risk, as well as the cell type of action. To this end, we utilized GWAS datasets for type 1 diabetes and type 2 diabetes, pancreatic cancer, BMI, WHR, WHRadjBMI, fasting glucose, fasting insulin, HBA1c, 2-hour glucose, total cholesterol, triglycerides, HDL, LDL, sleep duration, insomnia, and Alzheimer’s disease. We tested these categories against both our single cell eQTLs and deconvoluted eQTLs to gain insight on associations with both datasets.

For our single cell eQTLs, we found significant associations (PPH4 > 0.8) with eight eQTLs across four different eGenes. The eGene *ADCY5* had the most significant co-localization, with an eVariant found in alpha cells which co-localized with fasting insulin, fasting glucose, 2-hour glucose, HbA1c levels, and type 2 diabetes **(Supp Table 21)**. Variants in *ADCY5* have been frequently associated with type 2 diabetes risk[64] and were previously implicated in both the InsPIRE study[18] and TIGER study[17] **(Supp Table 6, Supp Table 7).** There were two other co-localizations with type 2 diabetes, *GALNT3* with an alpha cell variant and *PM20D1-AS1* with a beta cell variant **(Supp Table 21)**. As characterized earlier, *GALNT3* was a significant eQTL that we found only in alpha cells that was previously implicated in islet eQTLs from InsPIRE[18]. The last colocalization was rs2168101 and *LMO1* in alpha cells for fasting glucose. *LMO1* has been implicated in glucose levels previously[65] and the SNP-gene pair we identified is a known eQTL in skeletal muscle from GTEx and pancreatic islets from TIGER.

There were 65 significant co-localizations with the deconvoluted eQTLs, including 28 for acinar cells, 32 for ductal cells, 4 with beta cells, and 1 with alpha cells **(Supp Table 22)**. These co-localizations were mainly associated with type 2 diabetes, but also included hits from fasting glucose, 2-hour glucose, WHR, WHRadjBMI, BMI, HbA1c levels, Alzheimer’s disease, and sleep duration **(Supp Table 22)**. One connection identified was an eQTL for *LRRC37A* in acinar cells that co-localized with sleep duration. *LRRC37A* has been implicated in sleep traits previously as well as some metabolic traits, like CAD[66]. There is also emerging literature showing a potential connection between the pancreas and sleep[62]. Overall, between both the deconvoluted eQTLs and single cell eQTLs, we observed candidate disease related eQTLs that warrant follow-up.

### Association of rs2168101 and *LMO1*

One eQTL we followed up on due to its co-localization with a GWAS variant, was rs2168101 for *LMO1* in alpha cells in our single cell data. We found this variant was associated with fasting glucose level in GWAS from the MAGIC consortium[36] **(Supp Table 21)** and in fact was the sentinel variant at this locus **(Fig 4a)**. The alpha cell eQTL showed decreased expression for *LMO1* for rs2168101-A. **(Fig 4b)** and for MAGIC’s fasting glucose GWAS, rs2168101-A was associated with lower fasting glucose levels[36]. Interestingly, increased *LMO1* expression has been connected to several different cancers, including prostate cancer, non-small cell lung cancer, and neuroblastoma[67–9]. Additionally, in the case of neuroblastoma, rs2168101-A is associated with reduced risk, and also to decreased *LMO1* expression in several neuroblastoma cell lines[70]. We did not discern evidence of an eQTL effect for *LMO1* in other cell types **(Fig 4c)**. We also did not observe evidence of an eQTL effect in our deconvoluted eQTL analysis, consistent with the small number of eQTLs detected in deconvoluted alpha cells from GTEx. When compared to previous pancreatic studies, we did not find any significant eQTLs for *LMO1* in GTEx or InsPIRE but did find an eQTL for TIGER. However, it did not co-localize with our alpha cell eQTL **(Supp Table 5)**. When looking across other tissues, we found that rs2168101 associated with *LMO1* is an eQTL in GTEx skeletal muscle, also with a negative direction of effect[19].

Our fine mapping indicated rs2168101 as the only causal variant in the credible set, with a PIP value of > 0.99 **(Supp Table 11).** This limited the scope of other variants that could be followed up on but was consistent as there were no other variants achieving statistical significance for an alpha cell eQTL within this region **(Fig 4c).** When analyzing cell-type specific transposase-accessible chromatin data, we found that rs2168101 was in open chromatin in the endocrine cell types (alpha, beta, delta, and gamma), but not in the exocrine acinar or ductal cells **(Fig 4d).** This supports the hypothesis that rs2168101 is an important variant for study with respect to fasting glucose levels, especially in alpha cells, which are known to regulate glucose levels through their release of the hormone glucagon[5].

## DISCUSSION

In this study, we sought to gain a greater understanding of pancreatic biology at the single cell level leveraging single-cell pancreatic islet data from HPAP. We analyzed how variants influence gene expression on a cell type specific level through eQTL analysis. While previous studies have carried out eQTL discovery in the pancreas, this has not been previously performed at the single cell level. By executing discovery at the cell type level, we hypothesized that we could map known bulk eQTLs to the cell types that were most relevant and discover new eQTLs that went previously undetected in bulk studies. We leveraged the single cell data in multiple ways to accomplish this goal: first by using the single cell data itself to carry out eQTL discovery, then by using the single cell data as a reference to deconvolute bulk pancreatic data and pursue eQTL discovery on the deconvoluted bulk data.

eQTL discovery of the single cell data itself enabled us to identify 112 unique eGenes across 6 pancreatic cell types (alpha, beta, delta, gamma, acinar, ductal). Most of the eQTL and eGenes revealed were significant in just one of these cell types rather than shared across multiple. When compared to previous bulk pancreatic studies, we found that 99 of our eGenes were reported previously and 13 were novel discoveries. We determined the likely relevant cell type of action for multiple previously known eQTLs, such as *GALNT3* at rs1968293 in alpha cells.

When we moved to conducting discovery in deconvoluted eQTLs, we revealed a larger number of significant eQTLs, but in fewer cell types. Overall, we observed 3,134 unique deconvoluted eGenes across four pancreatic cell types (alpha, beta, acinar, ductal) with most of these significant eGenes being in acinar and ductal cells. Bulk eQTLs are mainly driven by these cell types, as they are the most abundant in the pancreas. When compared with previous studies, we found 116 of our significant eGenes were unique to our analysis, the majority of which were found in ductal cells. Unsurprisingly, a majority of our eGenes were also present in GTEx’s pancreatic eQTL study, as the bulk data we used were from GTEx.

Post-eQTL experiments such as fine-mapping and co-localization with disease GWAS results provided further insight into which eQTLs warrant future follow-up in the pancreas. One such locus is rs2168101 at *LMO1*, which was a significant eQTL in alpha cells that co-localized with the MAGIC consortium’s fasting glucose GWAS signal. This variant-gene pair is known to confer an effect across many cancers. A previous study that explored this variant to gene connection in neuroblastoma proposed that a disruption in GATA binding could be a mechanism for how rs2168101 effects *LMO1* expression[70]. The transcription factor GATA6 is known to be expressed in pancreatic endocrine cells[71] and *GATA6* is expressed in the alpha cell population of both the HPAP single-cell data and deconvoluted alpha cell population. Disruption in GATA binding might be affecting *LMO1* expression in alpha cells as well. Experimental studies will be necessary to test this hypothesized mechanism and to further explore rs2168101 as a variant that could affect how alpha cells regulate fasting glucose levels.

There are several limitations to our study, with the most obvious relating to discovery power. Firstly, we had to be stringent with our quality control measures for our single cell data in order to label cell types confidently, which could cause us to lose power or miss other interesting patterns within the data. Also, while utilizing single-cell data to carry out eQTL studies can rule in cell types of interest for eQTLs, due to the degree of data available for discovery, we could not rule out that a given eQTL might be implicated in more cell types if more cells and individuals had been analyzed. With only 71 samples at most, we are only powered to detect the strongest eQTLs. While utilizing deconvolution can make use of single-cell data to generate more powered data, there are significant limitations to this approach as well. With deconvolution, the gene expression profiles are imputed rather than measured experimentally, inevitably adding noise and uncertainty to the data. Beyond that, studies like GTEx that have RNA-seq data from the entire pancreas will overwhelmingly represent the more common cell types, such as the exocrine acinar and ductal cells that make up the majority of the pancreas. The relatively rare endocrine cells will therefore be underpowered for discovery as well. These limitations for deconvolution also highlight the importance of producing high powered single cell databases in the future for cell type specific studies. Another limitation of this study is that the donors represent multiple disease states across type 1 and type 2 diabetes as well as non-diabetic donors. Ideally, this study would be done in non-diabetic donors only to reduce uncertainty in transcript levels that are modulated in a disease state; however, excluding diabetic donors in this case would have made the power too low for eQTL discovery.

Overall, we present a resource for analyzing cell type specific pancreatic eQTLs using both scRNA-seq data and deconvoluted bulk pancreatic data. While single cell datasets can be limited in the number of individuals or cells sequenced currently, we also demonstrate that such datasets can successfully be utilized to leverage and deconvolute larger bulk datasets. In future studies, it will be important to assess more modalities at cell type resolution to understand pancreatic biology in more depth. As technology improves, consortia like HPAP can move to carry out more advanced approaches, such as multiomics, which will drive future additional discoveries.

## AKNOWLEDGEMENTS

We extend our gratitude to all donors, families and coordinators contributing to the Human Pancreas Analysis Program (HPAP; RRID:SCR_016202; PMID: 31127054; PMID: 36206763). The Genotype-Tissue Expression (GTEx) Project was supported by the Common Fund of the Office of the Director of the National Institutes of Health (commonfund.nih.gov/GTEx). Additional funds were provided by the NCI, NHGRI, NHLBI, NIDA, NIMH, and NINDS. Donors were enrolled at Biospecimen Source Sites funded by NCI\Leidos Biomedical Research, Inc. subcontracts to the National Disease Research Interchange (10XS170), Roswell Park Cancer Institute (10XS171), and Science Care, Inc. (X10S172). The Laboratory, Data Analysis, and Coordinating Center (LDACC) was funded through a contract (HHSN268201000029C) to the Broad Institute, Inc. Biorepository operations were funded through a Leidos Biomedical Research, Inc. subcontract to Van Andel Research Institute (10ST1035). Additional data repository and project management were provided by Leidos Biomedical Research, Inc.(HHSN261200800001E). The Brain Bank was supported supplements to University of Miami grant DA006227. Statistical Methods development grants were made to the University of Geneva (MH090941 & MH101814), the University of Chicago (MH090951,MH090937, MH101825, & MH101820), the University of North Carolina - Chapel Hill (MH090936), North Carolina State University (MH101819), Harvard University (MH090948), Stanford University (MH101782), Washington University (MH101810), and to the University of Pennsylvania (MH101822).

## DATA AVAILABILITY

scRNAseq data from the Human Pancreas Analysis Program (HPAP) is available though PANC-DB (https://hpap.pmacs.upenn.edu/). Whole genome sequencing from HPAP is available through dbGAP (accession no. phs002465.v2.p1). Gene reads per tissue from the Gene-Tissue Expression (GTEx) project are available from the GTEx Portal (https://gtexportal.org). Whole genome sequencing from GTEx is available through dbGAP (accession no. phs000424.v10.p2).

## AUTHOR’S RELATIONSHIPS AND ACTIVITIES

The authors have no conflicting interests to disclose.

## CONTRIBUTION STATEMENT

MAW, SFAG, BFV conceived of the study. MAW acquired the data. MAW, KL, RE analyzed the data. MAW, SFAG, and BFV prepared the original draft of the manuscript. SFAG and BFV supervised the project. All authors interpreted the data, reviewed critically, edited content, and approved the version to be published.

## ABBREVIATIONS

ABF: approximate bayes factor
eGene: gene associated with an eQTL
eQTL: expression quantitative trait loci
eVariant: variant associated with an eQTL
FDR: false discovery rate
GTEx: Genotype Tissue Expression project
GWAS: genome-wide association study
HPAP: Human Pancreas Analysis Program
PPH4: posterior probability of hypothesis four
scRNA-seq: single cell RNA sequencing
TPM: transcripts per million
UMI: unique molecular identifier
WGS: whole genome sequencing
WHR: waist-hip-ratio
WHRadjBMI: waist-hip-ratio adjusted for BMI

