## Supplemental Figures for "Utilizing single-cell data for per-cell type eQTL mapping in the human pancreas"

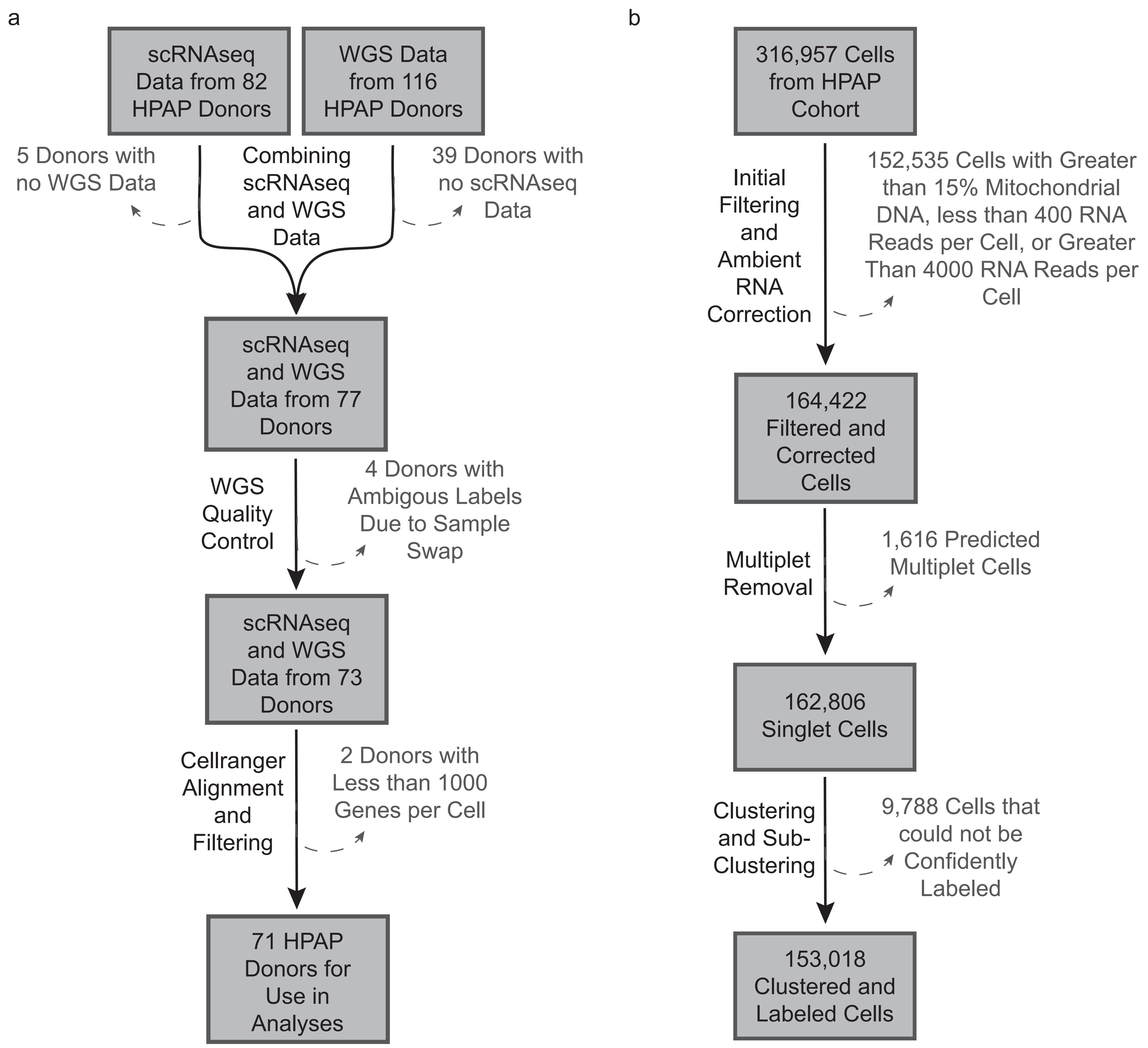


**Supplementary Figure 1 Initial Quality Control on Raw Single Cell Sequencing Data:** (a) Steps for filtering from initial donor pool to the 71 donors included in the final analysis. (b) Steps for filtering the cells from the 71 donors to the final number of cells analyzed.

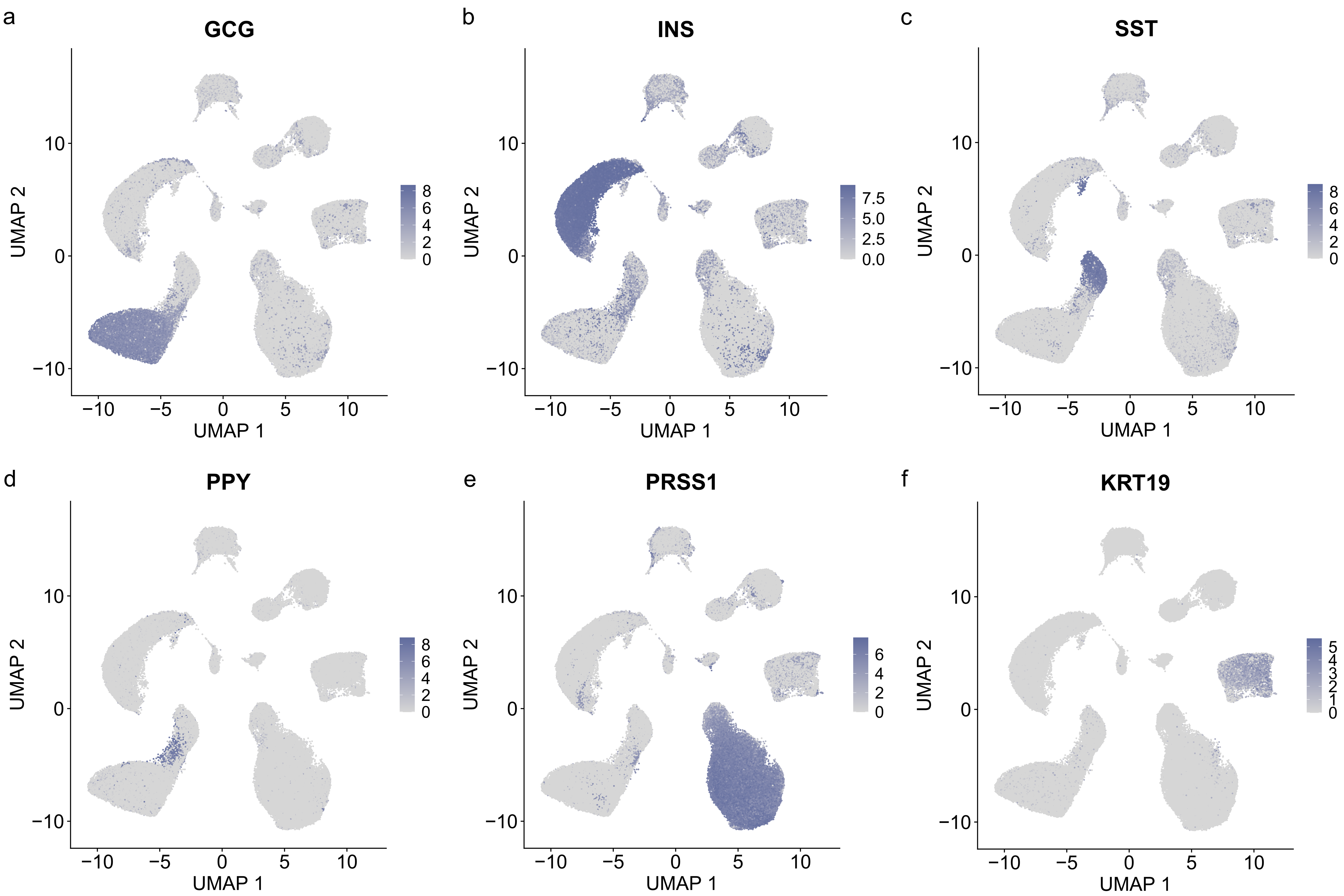


**Supplementary Figure 2 Distribution of Marker Genes across UMAP clustering:** Distribution of common pancreatic marker genes across UMAP clusters. Marker genes are for (a) insulin (beta cells), (b) glucagon (alpha cells), (c) somatostatin (delta cells), (d) pancreatic polypeptide (gamma cells), (e) protease serine 1 (acinar cells), (f) keratin 19 (ductal cells).

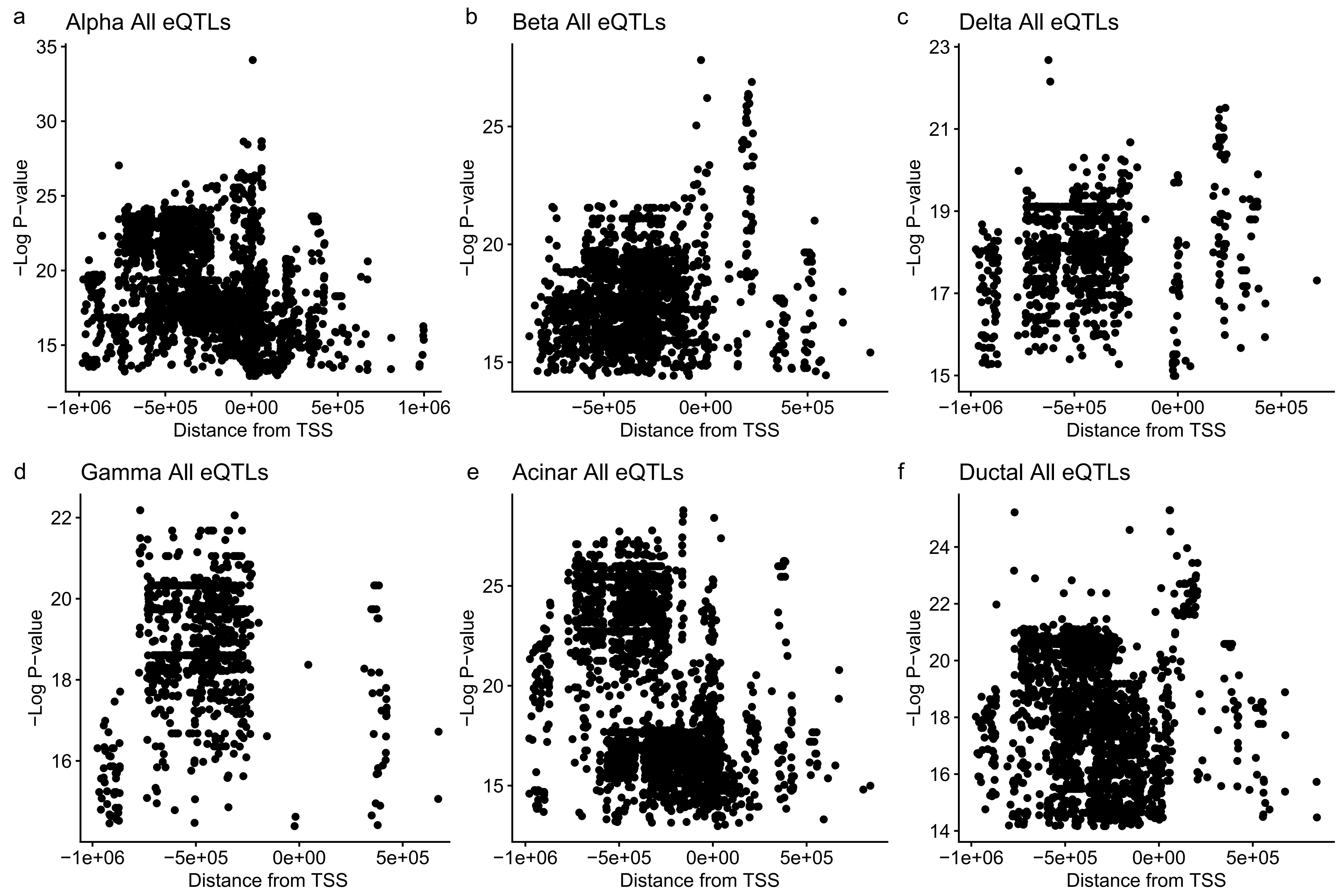


**Supplementary Figure 3 P-value of HPAP single cell eQTLs versus distance from transcription start site:** Graphs showing the p-value of eQTLs versus their eVariant's distance from the eGene's transcription start site. This was done per cell type showing (a) alpha, (b) beta, (c) delta, (d) gamma, (e) acinar, (f) ductal

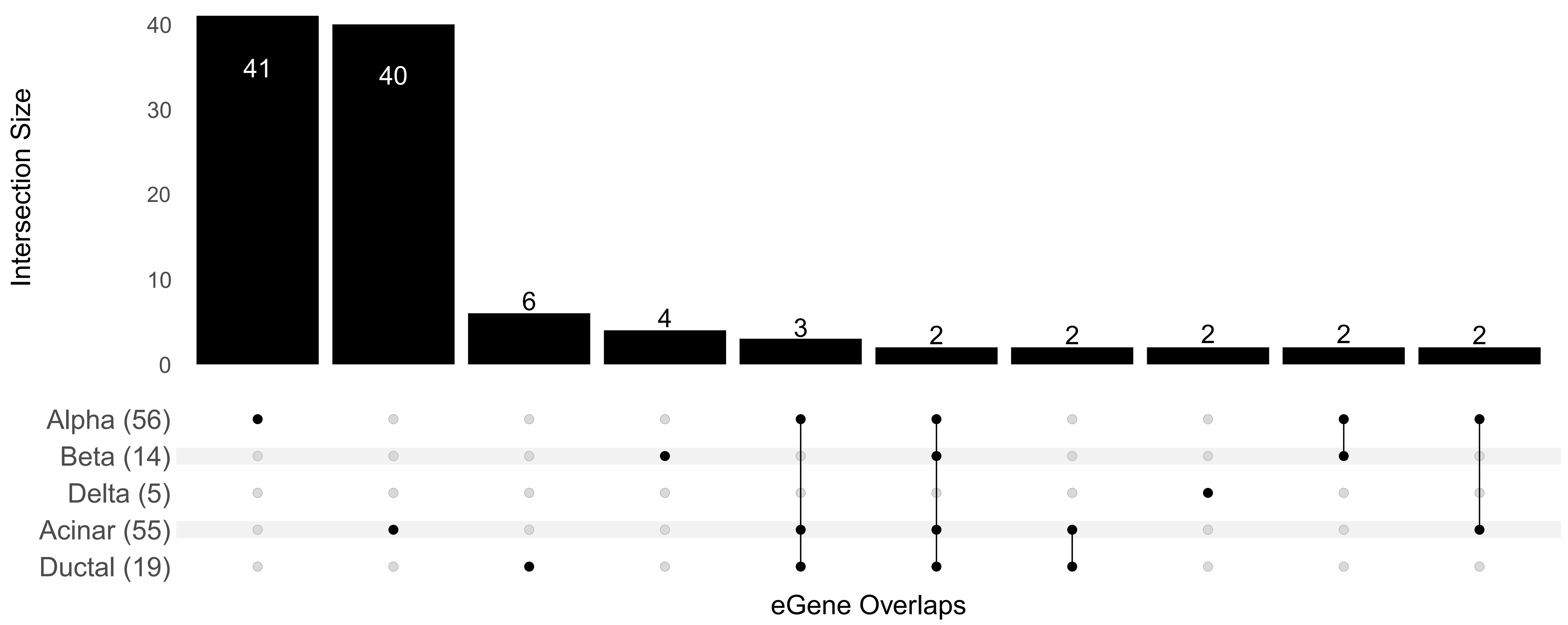


**Supplementary Figure 4 Overlap between HPAP single cell eQTL eGenes across cell types:** Upset plot showing overlap of eGenes between the different single cell eQTL discovery sets from HPAP.

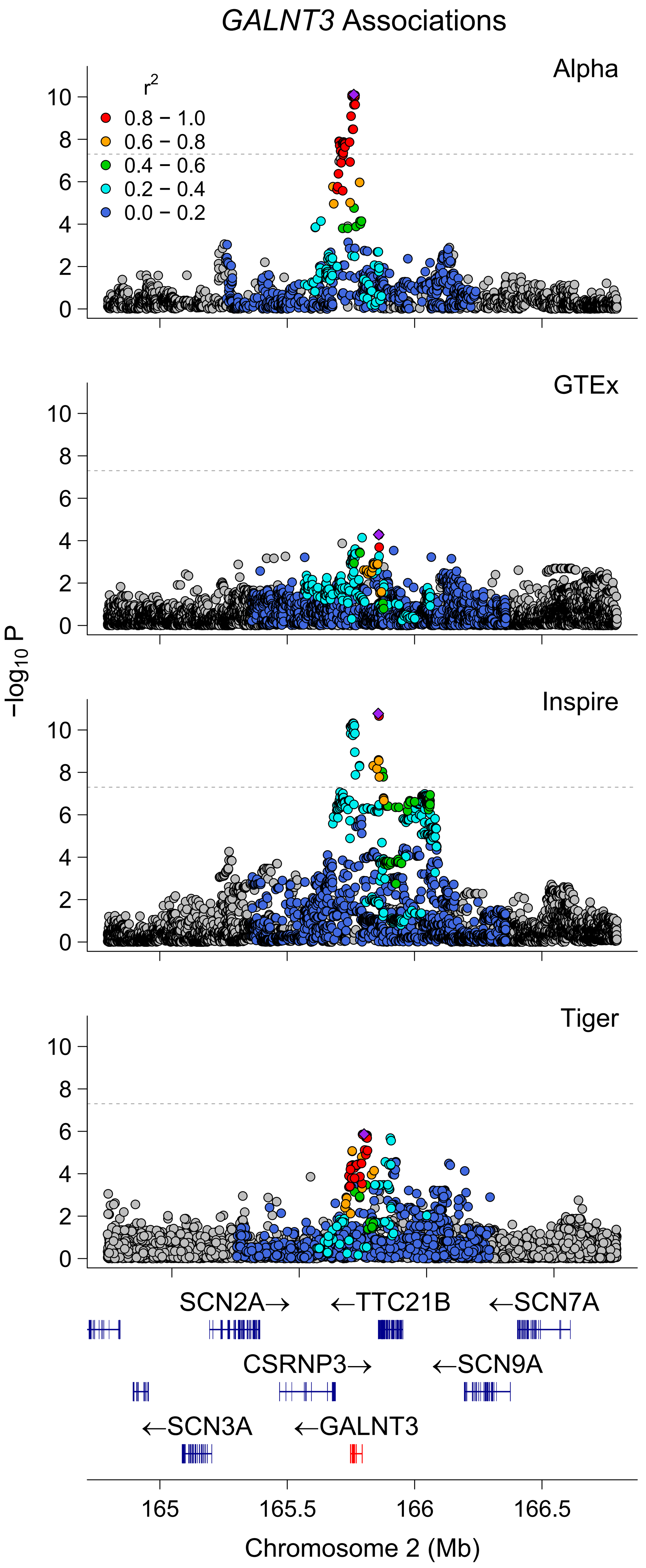


**Supplementary Figure 5 GALNT3 across previous studies:** Locus zoom plot of the GALNT3 locus across the HPAP single cell alpha eQTL study and three previous pancreatic eQTL studies (GTEx bulk pancreatic eQTL discovery, InsPIRE islet eQTLs, and TIGER islet eQTLs).

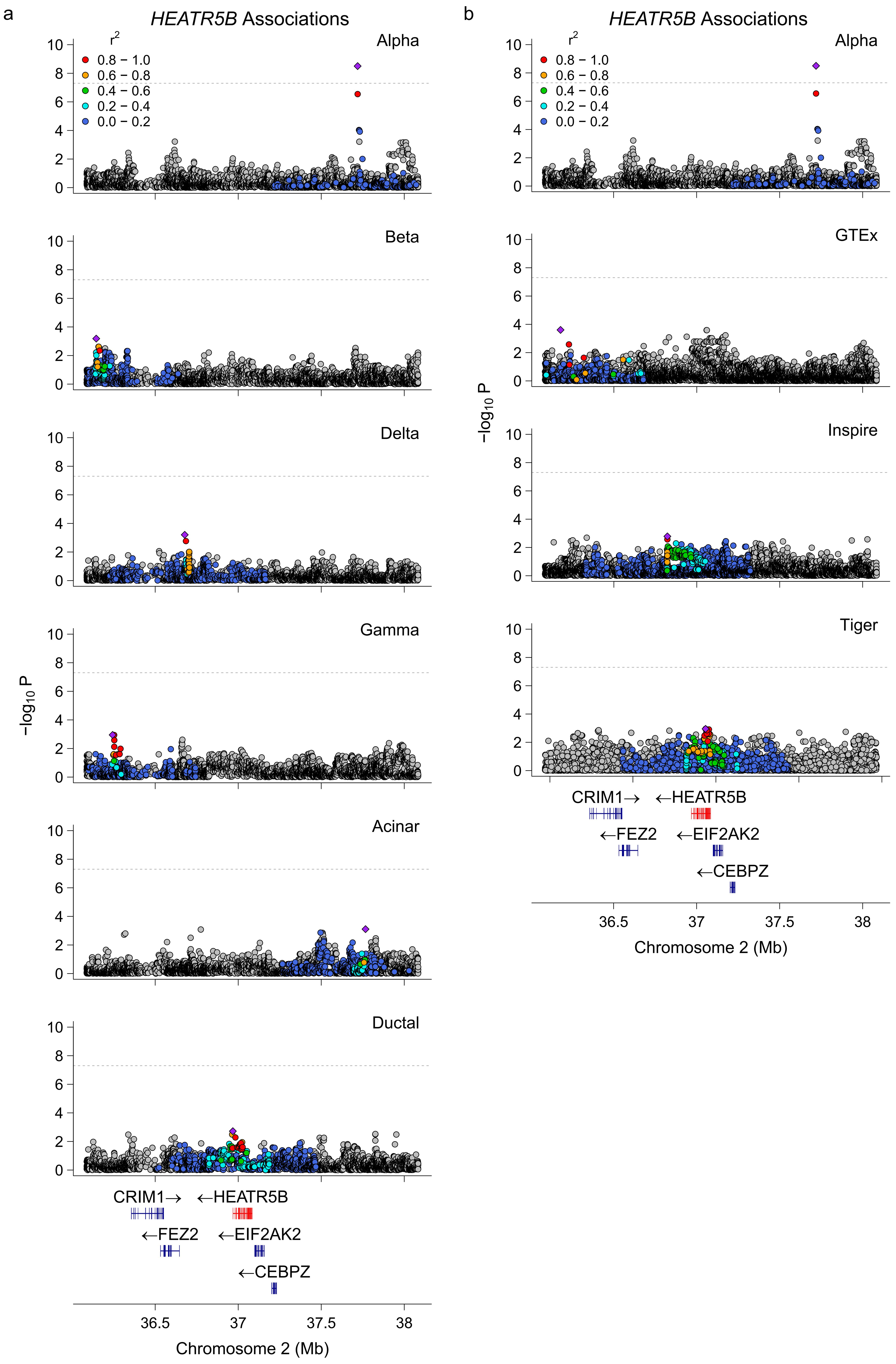


**Supplementary Figure 6 HEATR5B locus:** Locus zoom plot of the HEATR5B locus within (a) the HPAP single cell eQTL data and (b) The alpha cell HPAP single cell eQTL discovery and previous pancreatic eQTL studies (GTEx bulk pancreatic eQTL discovery, InsPIRE islet eQTLs, and TIGER islet eQTLs).

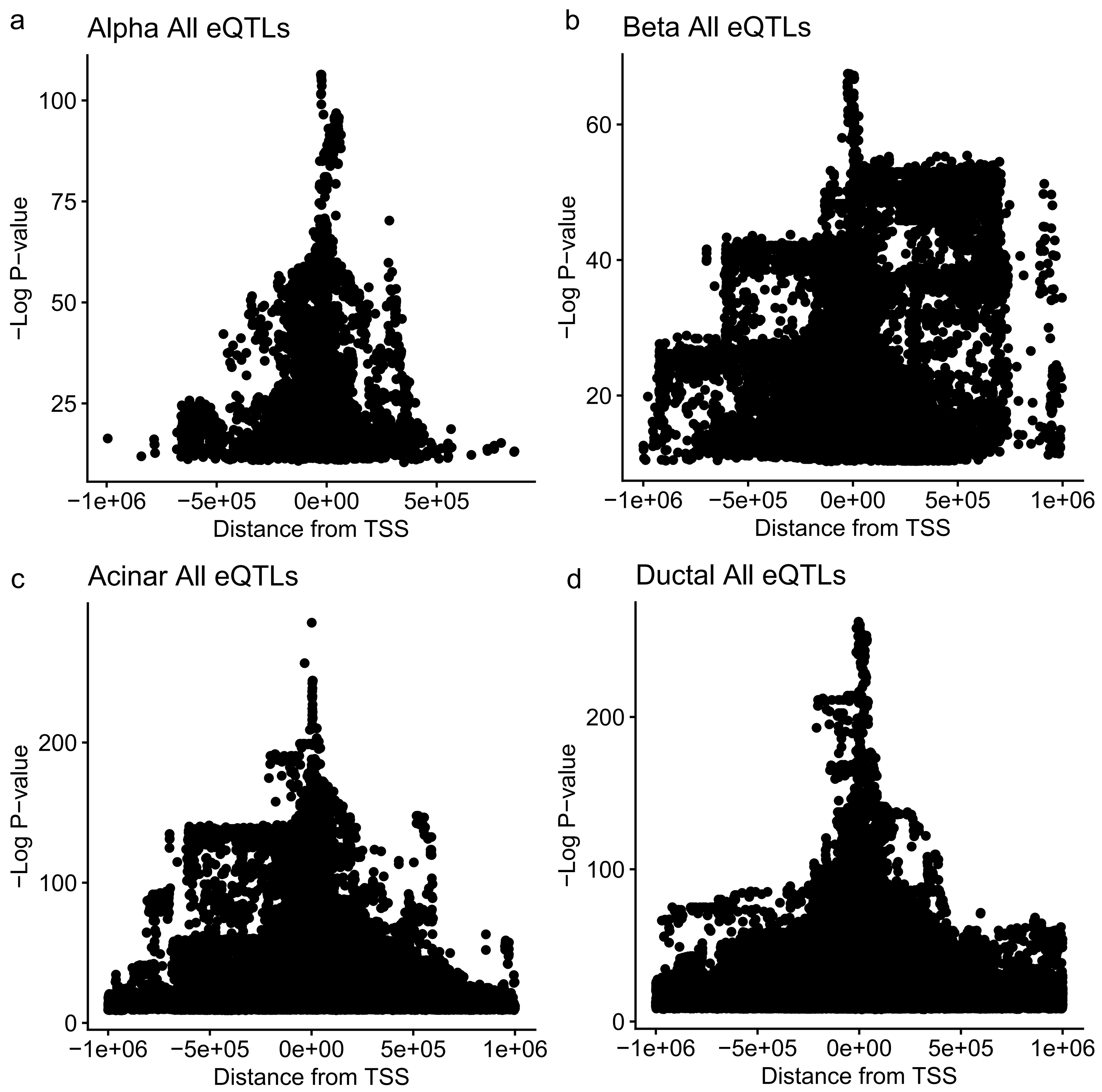


**Supplementary Figure 7 P-value of Deconvoluted eQTLs Versus Distance from Transcription Start Site:** Graphs showing the p-value of eQTLs versus their eVariant's distance from the eGene's transcription start site. This was done per cell type showing (a)alpha, (b) beta, (c) acinar, and (d) ductal

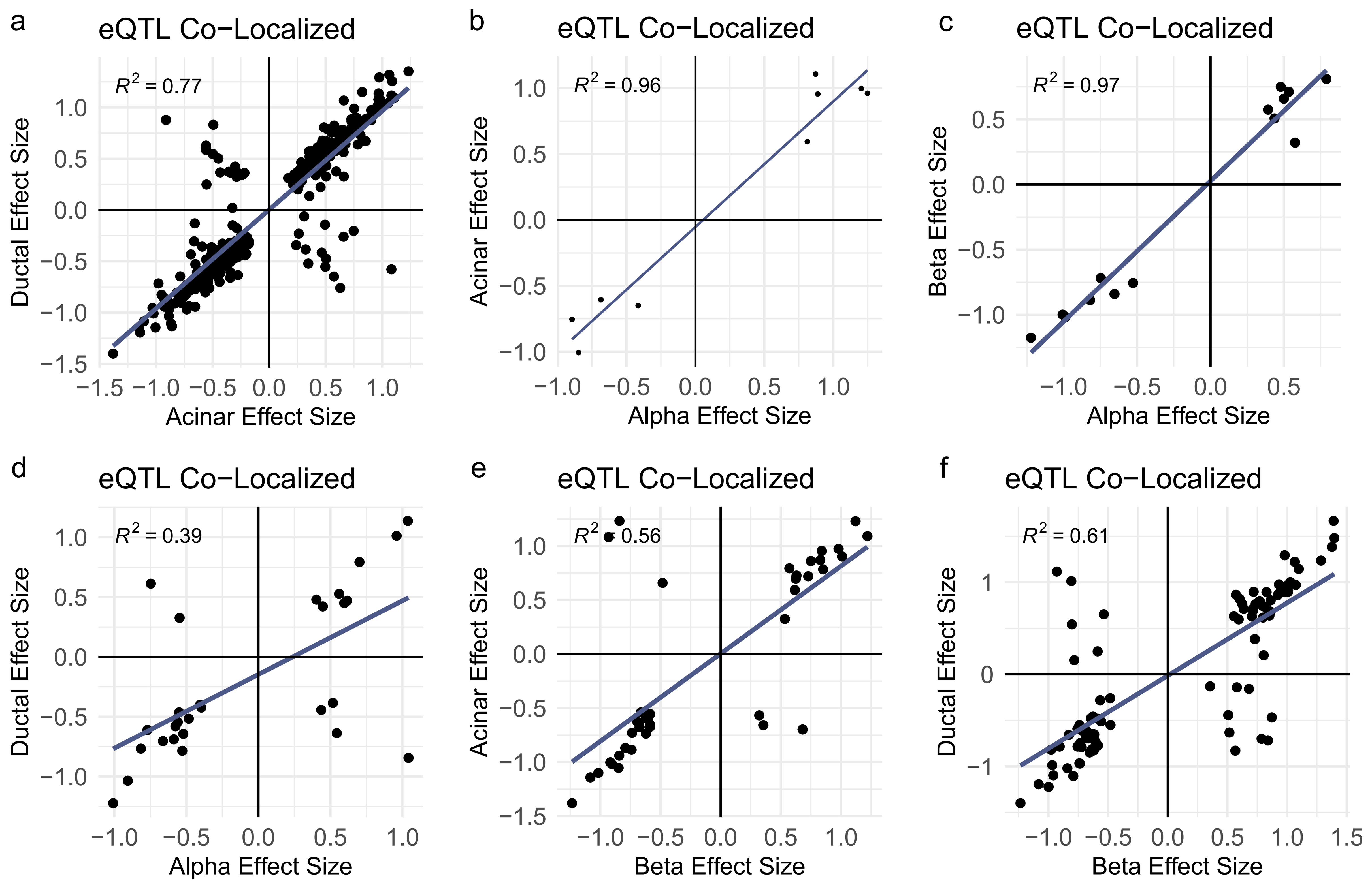


**Supplementary Figure 8 Direction of Effect Comparison Across Deconvoluted Cell Type eQTLs:** Plots showing the direction of effect compared for co-localized eQTLs between the different cell type deconvoluted eQTLs. (a) ductal and acinar cells, (b) acinar and alpha cells, (c) beta and alpha cells, (d) ductal and alpha cells, (e) acinar and beta cells, (f) ductal and beta cells.

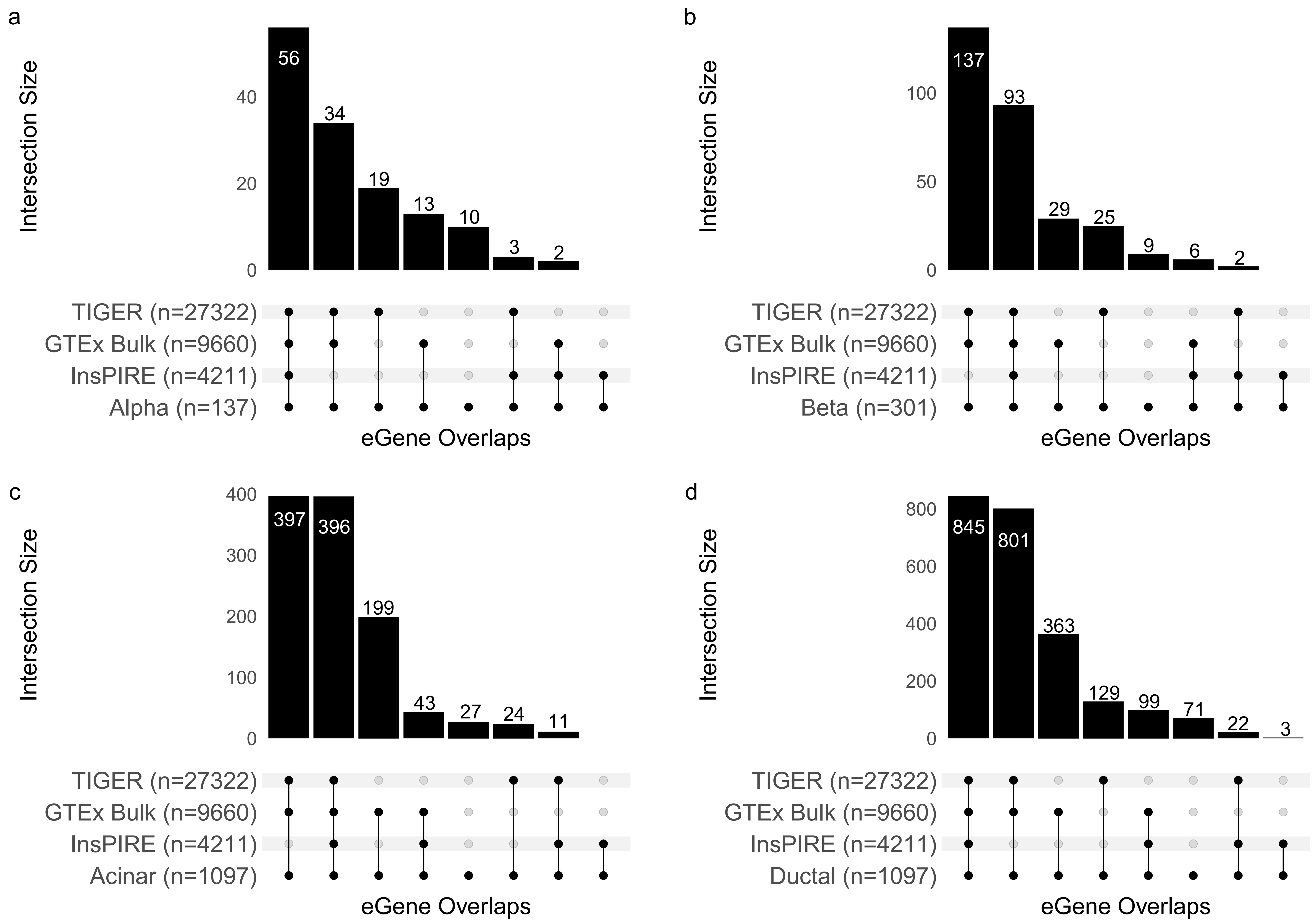


**Supplementary Figure 9 Per cell type eGene comparison between deconvoluted eQTLs and previous studies:** Upset plots showing the comparison between each deconvoluted cell type eQTLs and previous pancreatic eQTL studies. (a) alpha cells compared to previous studies, (b) beta cells compared to previous studies, (c) acinar cells compared to previous studies, (d) ductal cells compared to previous studies
